# Tissue tearing degrades optimal-transport and diffeomorphic registration of spatial transcriptomics beyond displacement magnitude: a multi-seed deformation benchmark and a supervised graph cross-attention proof-of-concept

**DOI:** 10.64898/2026.06.30.735390

**Authors:** Rushil Maniar, Sean Lee, Sunyoung S. Lee

## Abstract

**Background:** Three-dimensional reconstruction from serial spatial-transcriptomics (ST) sections requires registering adjacent slices, but physical sectioning introduces tears — discontinuous, non-isometric deformations. Leading methods rely on priors that tears strain: PASTE/PASTE2 use Fused Gromov–Wasserstein optimal transport (OT), which assumes near-isometric preservation of within-slice distances, while STalign and CODA use diffeomorphic (LDDMM) mapping, which cannot change tissue topology. Learned-deformation ST methods are emerging (STaCker, INST-Align), but OT/diffeomorphic behaviour under tearing has not been systematically characterised.

**Methods:** On the spatialLIBD human DLPFC Visium dataset (Maynard et al., 2021; 3 donors), we build a controlled benchmark — known smooth warps, single-block rigid tears (expression unchanged), and an identity self-control — at severities of 0–8 spot pitches, scored against an approximate array-position ground truth (∼8 px residual). We evaluate three unsupervised incumbents — PASTE2 (OT, over five warp seeds), STalign (diffeomorphic LDDMM), and GPSA (Gaussian-process warp) — add a magnitude-matched smooth control, and test a minimal graph model, Sutura (per-slice graph encoder →cross-attention correspondence → per-spot displacement; spatial coupling is local kNN message passing only, no explicit smoothness penalty). Sutura is trained supervised on each tissue’s ground truth; all baselines are unsupervised. Generalisation is assessed by leave-one-donor-out across all three donors.

**Results:** OT registration is robust to smooth warps but degrades reproducibly under tearing: nearest-correspondence (argmax) error 722 ± 5 →855 ± 27 px and layer accuracy 64.9% → 60.5% (mean ± 95% CI, 5 seeds). The effect is not merely displacement magnitude: at a matched mean displacement ( ∼2000 px), a smooth warp costs 769 px / 60.2% accuracy whereas a tear costs 863 px / 57.5% — an extra ∼100 px and ∼3 points attributable to the discontinuity. STalign (LDDMM) and GPSA (GP warp) both collapse at severe tears (866 px and 931 px respectively), confirming tear-collapse is field-wide across three independent method families. Trained and evaluated on the same donor, Sutura fits torn-tissue correspondence to a median 99 →106 px (5-seed), but under leave-one-donor-out is 1236 ±2→ 1584 ± 52 px — ∼1.8–3.6× worse than PASTE2 on every unseen donor. A contrastive correspondence loss halves the gap on two of three donors (to 816→949 and 749→ 826 px, ∼1.1–1.2× PASTE2 at worst-case tear) but is modest on the third and never surpasses PASTE2.

**Conclusion:** Tearing is a real, magnitude-controlled failure mode of all three incumbent method classes. A learned model fits it in-sample but donor-invariant generalisation remains open. The contrastive fix roughly halves the held-out gap on two of three donors and nears PASTE2 at worst-case tear, but does not surpass it: donor-invariance is improved, not solved. The durable contribution is the benchmark, the characterisation across three method families, and an honest negative with a diagnosed mechanism.

## 1 Introduction

Spatial transcriptomics (ST) measures genome-wide expression while preserving each cell’s position in intact tissue, and was named Nature Methods’ Method of the Year in 2020. Annual publications have grown from under 100 to roughly 630 across more than 2,000 institutions worldwide. A central application is three-dimensional reconstruction: stacking serial tissue sections into a coherent volume to trace how transcriptional programs vary through an organ, tumour, or developing structure. That reconstruction is only as good as the registration step that aligns each section to its neighbour — and registration is harder than it looks.

Physical sectioning is destructive. As a microtome cuts thin sections, tissue routinely tears — a region detaches and shifts, folds back, or separates along a seam. Unlike the smooth stretching that mounting introduces, a tear is a discontinuous, non-isometric deformation: distances across the seam change abruptly and local topology is broken. Tears are common, visually obvious to anyone who has handled histology slides, and they corrupt exactly the cross-section correspondences that 3D reconstruction depends on. Despite their prevalence, tears have not been systematically studied as a failure mode for computational alignment methods.

The leading alignment methods rest on smoothness priors that tears strain. PASTE and PASTE2 [1, 2] cast registration as Fused Gromov– Wasserstein optimal transport (OT), which assumes near-isometric preservation of within-slice pairwise distances; a tear inflates the distortion cost of the true correspondence, so the optimum is pulled toward a wrong but distance-preserving match, and entropic regularisation further blurs the plan. STalign [7] and CODA [24] use diffeomorphic (LDDMM) mappings, which are smooth and invertible and therefore structurally cannot change topology — they cannot represent a tear at all. GPSA [6] learns a globally smooth Gaussian-process warp, which enforces continuity by construction. INST-Align [17] uses implicit neural representations with Jacobian regularisation enforcing smooth, volume-preserving deformations — the same structural limitation in a different mathematical form. None of these failure modes has been systematically characterised under controlled conditions.

We build a controlled tear benchmark on the human DLPFC Visium dataset [21] (3 donors): known smooth warps, single-block rigid tears, and an identity self-control across severities of 0–8 spot pitches, scored against an array-position ground truth. On it we quantify how three unsupervised incumbents — PASTE2, STalign, and GPSA — degrade under tearing versus matched-magnitude smooth warps. We then test a minimal supervised graph model (Sutura: per-slice graph encoder, cross-attention correspondence, per-spot displacement) in-sample and under leave-one-donor-out, and evaluate a targeted contrastive correspondence loss as a candidate fix for the diagnosed generalisation failure.

Our contribution is an honest one. The benchmark cleanly establishes that tearing is a real, magnitude-controlled failure mode across three independent method families: OT, diffeomorphic, and GP warp. The learned model fits torn tissue well in-sample but does not generalise across donors. A contrastive correspondence loss roughly halves the cross-donor gap on two of three donors yet never surpasses the unsupervised baseline out-of-sample. The durable result is the benchmark and the characterisation; donor-invariant learned registration remains open and is positioned as an explicit, diagnosed research problem rather than a solved one.

## 2 Results

### 2.1 Three independent method families all collapse at severe tears

On smooth warps PASTE2 is stable: barycentric median error is ∼647 px, flat across severity, and layer-transfer accuracy holds at ∼65% against an 18.7% random floor. Under tearing, nearest-correspondence (argmax) error rises from 722 ± 5 to 855 ± 27 px and accuracy falls from 64.9% to 60.5% (mean ± 95% CI, 5 seeds). The diffeomorphic baseline STalign is excellent on small deformation — 79 px median at severity 0 — yet collapses under the tear to 866 px, an ∼11× rise. GPSA, a learned Gaussian-process warp, is near-perfect at severity 0 (8.9 px) yet collapses identically to 931 px at severe tear. All three unsupervised method families — OT, diffeomorphic, and GP warp — converge to ∼840–930 px at severe tear, confirming tear-collapse is a field-wide failure mode rather than a limitation of any single approach (Figure 1).

**Figure 1:**
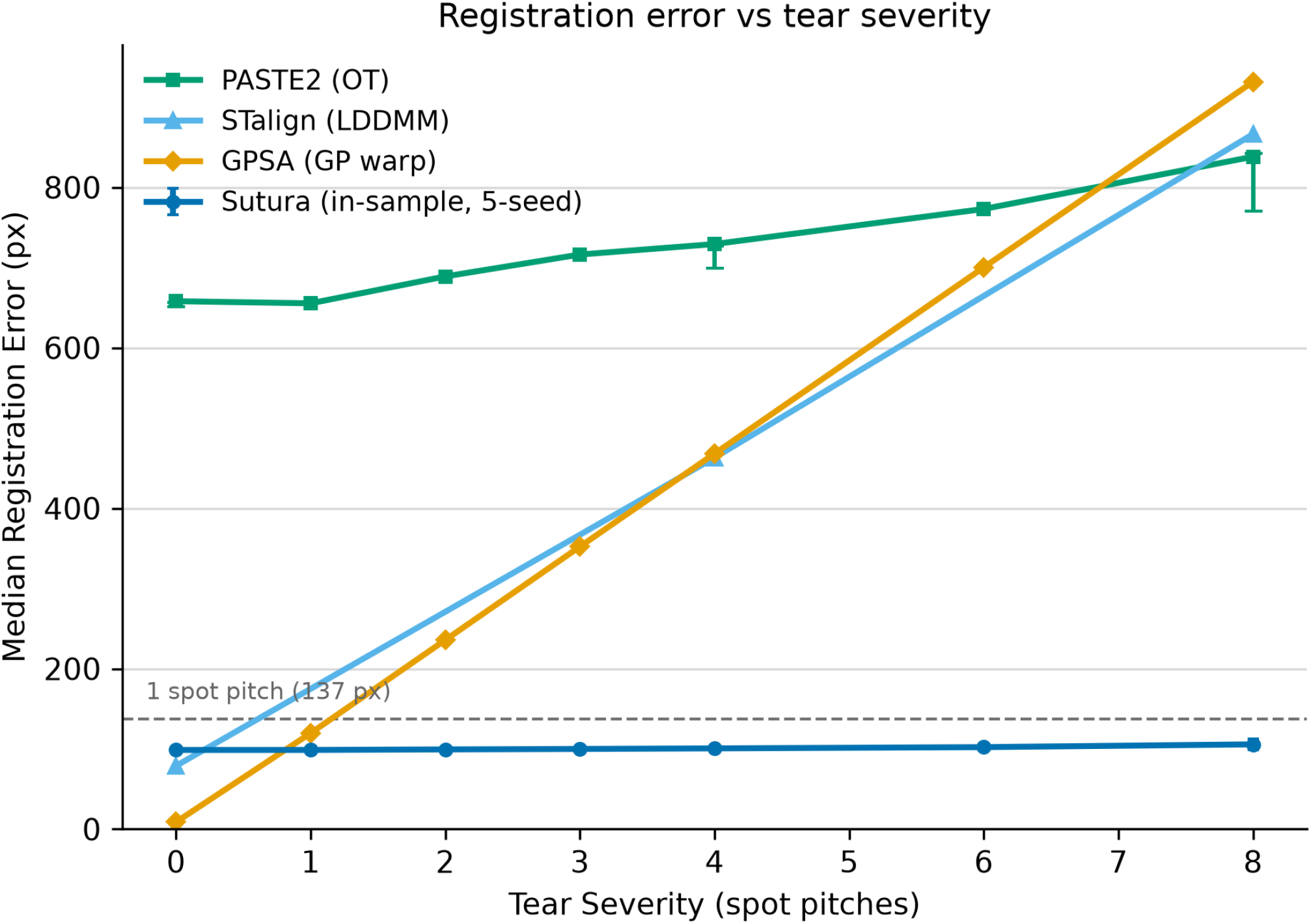
Registration error vs tear severity across all method classes. PASTE2 (OT), STalign (LDDMM), and GPSA (GP warp) all collapse at severe tears, converging to ∼840–930 px. Sutura (in-sample, 5-seed) remains flat at ∼106 px. Dashed line indicates 1 spot pitch (137 px). Error bars are 95% CI over 5 seeds where available.

### 2.2 The tear effect is not explained by displacement magnitude

Because the tear arm carries larger total displacement than the smooth arm at the same severity, we ran a magnitude-matched control: at equal mean displacement ( ∼2000 px), a smooth warp costs 769 px / 60.2% accuracy whereas a tear costs 863 px / 57.5% — an extra ∼100 px and ∼3 accuracy points attributable to the discontinuity itself, not its magnitude. The self-control registers at 0.0 px at every severity, validating the scoring pipeline (Figure 2).

**Figure 2:**
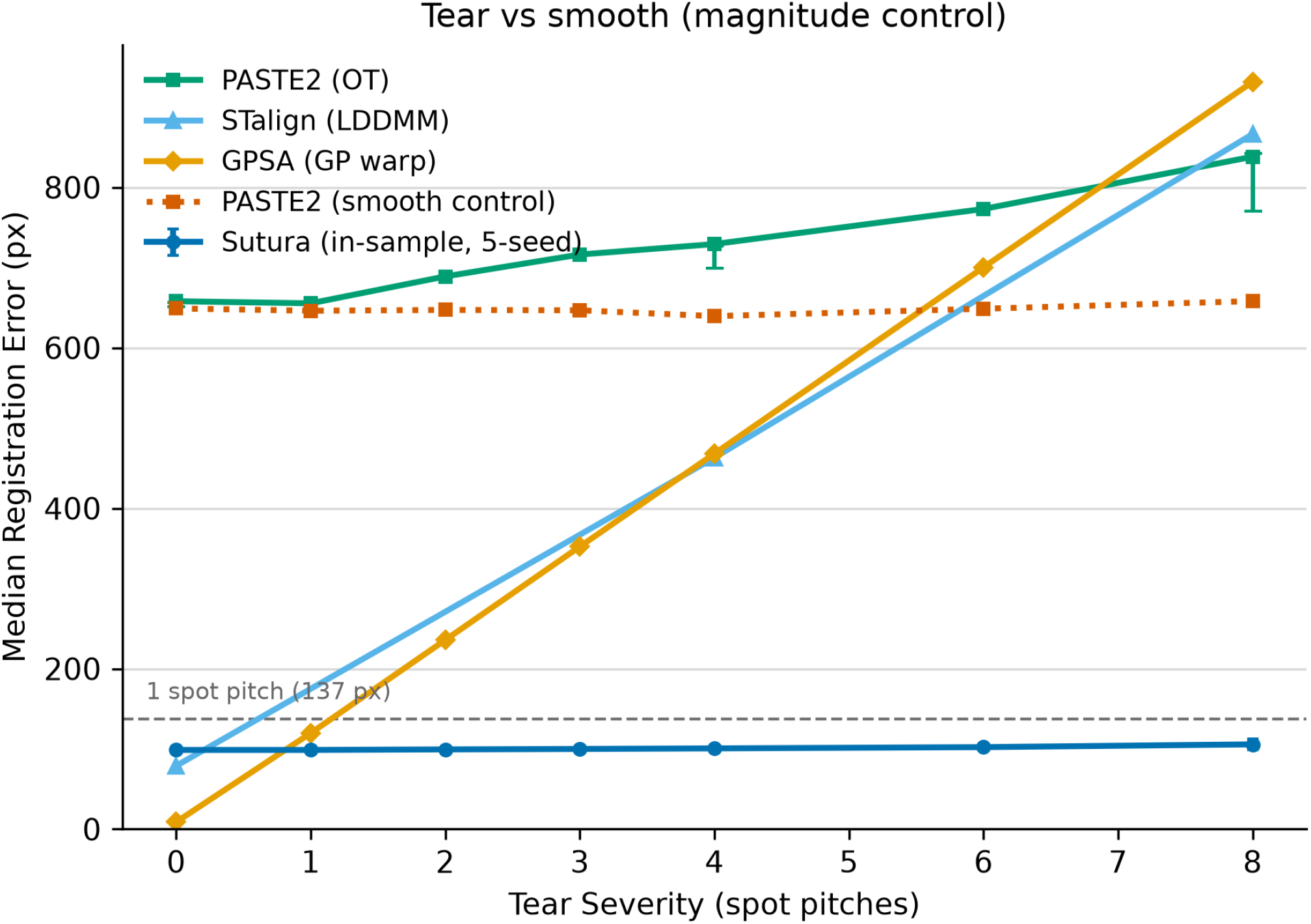
Magnitude-matched control. PASTE2 on smooth warps (dotted) stays flat while PASTE2 on tears rises, confirming ∼100 px of error is attributable to the discontinuity itself rather than displacement magnitude. STalign and GPSA collapse identically.

### 2.3 Sutura fits torn tissue in-sample, with caveats

Trained and evaluated on the same donor (151507/151508) with heldout warp seeds, Sutura fits torn-tissue correspondence to a median 99 → 106 px across severities (5-seed; an earlier single-seed report of 118 px was an unlucky draw). This is genuinely non-trivial against an expression-nearest-neighbour baseline of 2532 px, and a coordinate-shuffle control confirms the model uses geometry rather than ignoring it. Three honest caveats bound this result. First, Sutura is supervised on the exact ground-truth coordinates it is scored against, whereas all baselines solve cold. Second, forced to a hard one-spot-per-spot assignment, error is ∼138 px (*≈* 1 spot pitch) — part of the sub-pitch headline is interpolation between array positions. Third, the flat severity curve is partly structural: Sutura keys off local graph neighbourhoods and expression, both largely preserved by our synthetic warps.

### 2.4 The in-sample advantage does not survive across donors

Under leave-one-donor-out, Sutura’s advantage vanishes on all three donors. Held-out median error is approximately flat across severity but large: 1236±2→1373±34 px (per-slice standardisation) and 1429±3→1584±52 px (pooled) over five seeds on folds S2/S3, with S1 consistent at single seed — ∼1.8–3.6× worse than PASTE2 on the same unseen tissue (held-out PASTE2 397 ± 7 → 697± 18 px on S2; 528 ±7 → 539± 19 px on S3). The mechanism is diagnosed: cross-donor batch shift pushes held-out expression off-distribution in the training-fit SVD basis; the encoder produces non-discriminative embeddings; cross-attention collapses toward uniform; and predictions regress to the tissue centroid, producing a flat ∼1300–1600 px error across severity.

### 2.5 A contrastive correspondence loss narrows but does not close the gap

Adding an InfoNCE-style contrastive correspondence loss (single-seed, LODO on all three donors) cuts the held-out ga ∼38–42% on two donors: best held-out median falls to 816 → 949 px (S1) and 749 →826 px (S2), reaching ∼1.1–1.2× of PASTE2 at worst-case tear. The fix helps only modestly on the third donor: S3 improves from ∼1433 →1584 px to 1248 →1333 px ( ∼13%, still ∼2.4× PASTE2). The contrastive loss does not surpass PASTE2 on any donor. Donor-invariance is improved, not solved (Figure 3).

**Figure 3:**
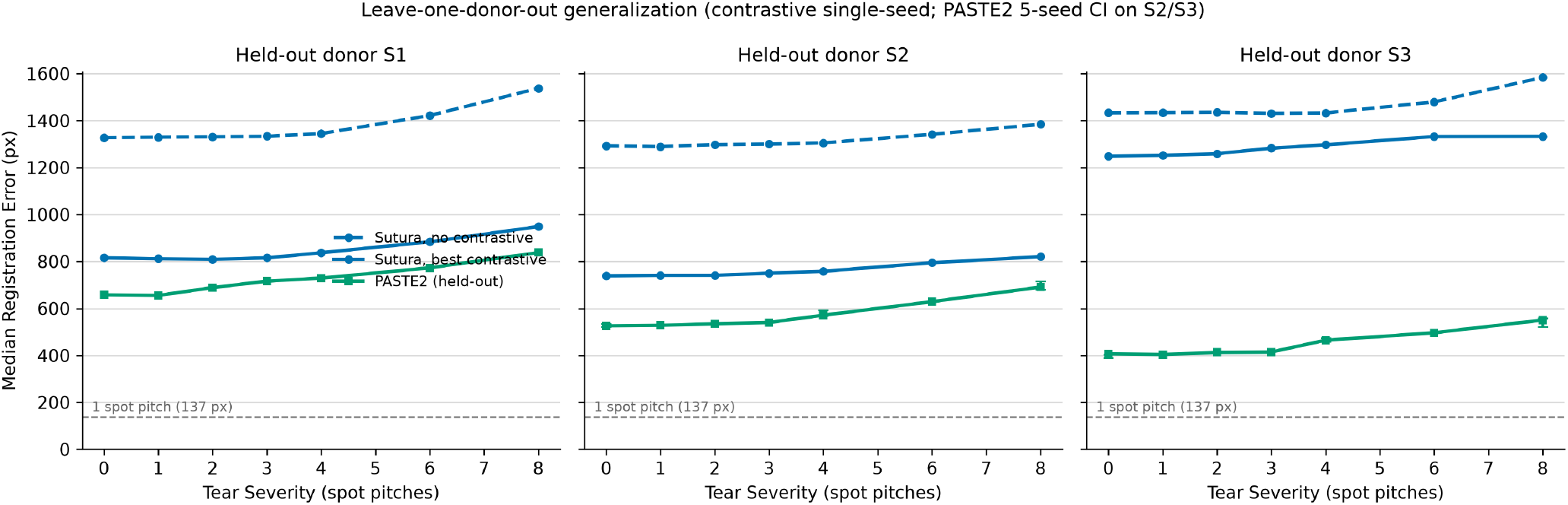
Leave-one-donor-out generalization. Across all three held-out donors, Sutura without contrastive loss (dashed) is 1.8–3.6× worse than PASTE2. The contrastive fix (solid) halves the gap on S1/S2 but helps only modestly on S3. PASTE2 5-seed CIs shown on S2/S3. Contrastive results are single-seed.

**Figure 4:**
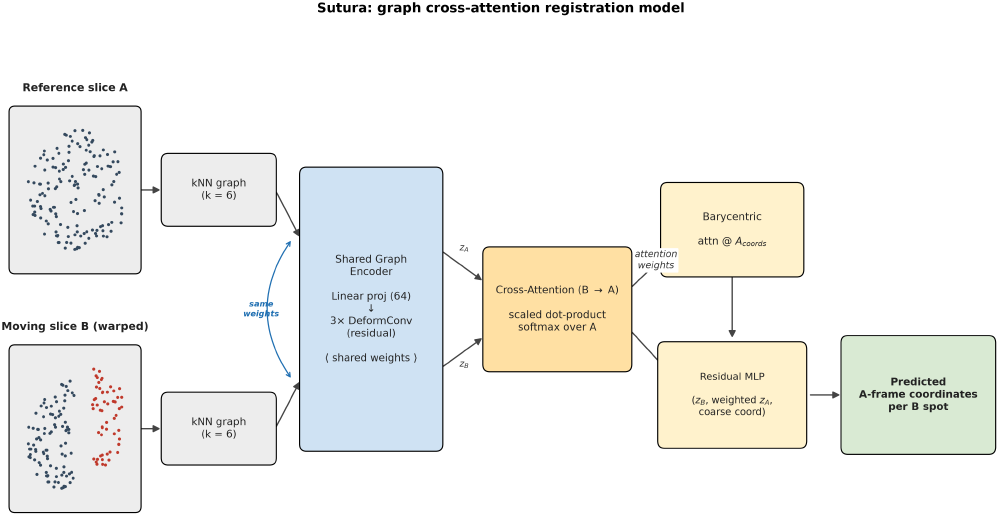
Sutura model architecture. Per-slice kNN graphs (*k* = 6) are encoded by a shared graph encoder (linear projection to width 64 followed by three residual DeformConv layers); cross-attention from moving slice (B) spots to reference slice (A) spots produces soft correspondences; predicted A-frame coordinates are the barycentric combination attn@*A*_coords_ plus a learned residual MLP correction.

## 3 Methods

### 3.1 Dataset

We used the human dorsolateral prefrontal cortex (DLPFC) 10x Visium dataset from the spatialLIBD resource [21], obtained as preprocessed .h5ad files from the CellCharter benchmark distribution (Figshare article 22004273, CC BY 4.0). We analysed three donors, each represented by two adjacent serial sections: 151507/151508 (S1, donor Br5292), 151669/151670 (S2, donor Br5595), and 151673/151674 (S3, donor Br8100). Each slice retains its full gene panel (33,538 genes) and carries manual cortical-layer annotations (L1–L6 plus white matter) in `obs[“layer”];` missing annotations are encoded as `“NA”`. Spot counts are 4,226/4,384 (S1), 3,661/3,498 (S2), and 3,639/3,673 (S3). The expression matrix (.X) was used only for node features and label-transfer scoring and was never modified during deformation. Within each donor pair, one section served as the fixed reference (A) and the adjacent section as the moving (B) slice. The spot pitch was defined per slice as the median nearest-neighbour distance of the spot centroids (∼137 px for the primary S1 pair).

### 3.2 Synthetic deformation benchmark

We applied known, parameterised deformations to the moving slice’s spatial coordinates `(obsm[“spatial”])` while leaving the expression matrix .`X` entirely unchanged. Two deformation classes plus an identity control were used, indexed by a dimensionless severity in spot pitches.

#### Smooth warp

A near-isometric displacement field was generated as the sum of *n*_bumps_ = 8 Gaussian radial bumps with random centres, directions, and amplitudes drawn uniformly from [0.5, 1.0], and a common width of 0.18× the bounding-box diagonal. The summed field was normalised to unit peak magnitude and scaled to a maximum displacement of severity × pitch.

#### Tear

A random cut axis (angle *θ ∼* Uniform[0, 2*π*)) was chosen; spots on one side of a threshold (quantile ∼ Uniform[0.55, 0.70], selecting ∼30–45% of tissue) were translated rigidly along the inplane perpendicular direction by 3× max(severity, 1) pitches, with the smooth field added on top. The expression matrix was left unchanged.

The standard severity grid was {0, 1, 2, 3, 4, 6, 8} spot pitches. Multi-seed evaluation used seeds {0, 9999, 10000, 10001, 10002}, disjoint from training seeds [1, 9998].

### 3.3 Ground truth (array bridge)

Adjacent Visium capture areas share array row/column indices, so a spot in B has a known correspondence to the A spot at the same `(array_row, array_col)`. Ground-truth A-frame targets were the coordinates of the A spot at B’s array position; B spots without an A match were masked. This array bridge matched *≈*95% of B spots in-sample and 95.3–96.1% on held-out folds. The residual floor is ∼8.1 px; hard one-spot assignment is quantised to ∼1 pitch ( ∼ 137 px).

### 3.4 Baselines

#### PASTE2 [2]

Partial Fused Gromov–Wasserstein OT; unsupervised. We ran `partial_pairwise_align` with *s* = 0.99, *α* = 0.1, and the glmpca dissimilarity. The transport plan was projected via barycentric (soft) and argmax (hard) projections.

#### STalign [7]

LDDMM diffeomorphism; unsupervised. Reference and moving slices were rasterised by spot density at *dx* = pitch*/*2 ( ∼68 px) and aligned by LDDMM with smoothness scale *a* = 5*· dx* and `niter`= 2000. Applied both in-sample and zero-shot to held-out donors.

#### GPSA [6]

Gaussian-process spatial alignment; unsupervised. Installed from `andrewcharlesjones/spatial-alignment` and applied to the identical benchmark. Validated at sev0 = 8.9 px confirming genuine alignment before the full sweep.

STaCker [16] (image-based; no public code) and INST-Align [17] (code unreleased) could not be run; no head-to-head numbers were fabricated. CODA [24] uses the same global-affine + LDDMM framework as STalign; STalign represents this diffeomorphic class.

### 3.5 Sutura model

Sutura is a supervised graph cross-attention proof-of-concept ( ∼74k parameters). Node features were a 50-dimensional TruncatedSVD embedding of library-size-normalised, log1p-transformed counts; the SVD basis was fit on training slices only. Per-slice kNN spatial graphs (*k* = 6, symmetrised) were built on each slice’s own coordinates with pitch-normalised relative-position edge features.

A shared graph encoder — a linear projection to hidden width 64 followed by three residual DeformConv message-passing layers — produced per-spot embeddings *z*_*A*_ and *z*_*B*_ . Cross-attention from each B spot to all A spots (scaled dot-product, softmax over A) gave attention weights; the predicted location was a barycentric combination attn@*A*_coords_ plus a learned residual MLP. The loss is plain mean L2 distance over array-bridge-matched spots. No smoothness or regularisation term was included.

### 3.6 Training

Each training step drew a fresh warp of B at severity sampled uniformly on [0, 8], a training-only seed, and tear probability 0.5. Optimisation used Adam (lr 10^*−*3^). The in-sample model was trained for 60 epochs × 8 steps; LODO models for 100 epochs × 12 steps.

### 3.7 Leave-one-donor-out

One SVD basis was fit on training donors only and applied to the held-out donor. We compared global standardisation (held-out donor standardised by training donors’ pooled mean/SD) and per-slice standardisation (each slice standardised by its own mean/SD). No external batch-integration method was applied.

### 3.8 Contrastive correspondence loss

An InfoNCE-style contrastive loss was added to sharpen cross-slice attention and resist collapse toward uniform weights. We tested two readout variants (cosine similarity, attention logits) and three loss weights (*λ* ∈ {0, 0.5, 2.0} ) under LODO on all three folds, single-seed. Temperature was fixed at 0.07.

### 3.9 Evaluation and statistics

Primary metric: Euclidean registration error in pixels between predicted and array-bridge ground-truth coordinates, reported as median, mean, and 90th percentile. Headline curves evaluated over five warp seeds, reported as mean ±95% normal-approximation CI across seeds. All randomness was seeded for reproducibility; computation was CPU-only.

## 4 Related Work

### 4.1 Slice alignment via optimal transport

The dominant approach casts ST alignment as optimal transport between spot sets. PASTE [1] aligns slices by fused Gromov– Wasserstein (FGW) OT, combining an expression-similarity term with a Gromov–Wasserstein term that matches within-slice pairwise distances. PASTE2 [2] extends this to partial FGW for partially overlapping tissue. Both inherit a near-isometry prior from the GW term, and entropic regularisation [4, 5] further diffuses the optimal plan. GPSA [6] learns a shared coordinate system via Gaussian-process deformation, also assuming smooth correspondence. Our benchmark targets exactly the regime where these priors are weakest: physical tears, where within-slice distances across the seam are not preserved.

### 4.2 Diffeomorphic registration

STalign [7] adapts LDDMM [8] to ST by rasterising spot density and computing a diffeomorphism (affine plus velocity field). By construction a diffeomorphism cannot change topology — it cannot represent the discontinuity a tear introduces. We confirm this empirically: STalign is near-perfect on small deformations ( ∼79 px median) yet collapses at the tear ( ∼866 px, ∼11× rise). CODA [24] applies the same global-affine + LDDMM framework; we take our STalign result as the representative benchmark for this diffeomorphic method class.

### 4.3 Learned-deformation and graph models

Several methods use spatial graphs and learned embeddings: STAligner [9], SLAT [10], CAST [11], SANTO [12], and GraphST [13]. Sutura sits in this lineage and is closest in spirit to correspondence-based registration from medical imaging (VoxelMorph [14], XMorpher [15]). We do not claim architectural novelty; Sutura is a deliberately minimal supervised proof-of-concept.

### 4.4 Recent alignment frameworks and the tear gap

INST-Align [17] achieves state-of-the-art OT Accuracy (0.702) and NN Accuracy (0.719) across nine datasets via implicit neural representations and Jacobian regularisation. Critically, Jacobian regularisation enforces smooth, volume-preserving deformations — the same structural limitation as LDDMM. INST-Align’s benchmarks cover smooth and large deformations only; torn tissue is not evaluated. STaCker [16] requires H&E histology and has no public code. Across all recent frameworks, the tissue tear regime remains unevaluated. Our benchmark is the first systematic characterisation.

### 4.5 Batch effects and cross-donor integration

Sutura’s diagnosed failure mode connects to ST batch-integration work — Harmony [18] and Scanorama [19] — none of which we apply, leaving donor-invariant correspondence as an explicit open problem.

## 5 Discussion

### 5.1 What the benchmark establishes

The primary contribution is a controlled characterisation of how three dominant unsupervised ST alignment paradigms fail under tissue tears. Against a shared identity self-control (0.0 px at every severity), two findings hold. First, all three method families — OT (PASTE2), diffeomorphic (STalign), and GP warp (GPSA) — collapse at severe tears, converging to ∼840–930 px by different mechanisms: OT smears its transport plan, the diffeomorphism overshoots smoothly across a seam it cannot represent, and the GP warp averages over the discontinuity. Second, this collapse is not explained by displacement magnitude: at matched mean displacement, a tear costs ∼100 px and ∼3 accuracy points more than a smooth warp.

### 5.2 The supervised proof-of-concept and its honest limits

Trained and evaluated on the same donor with held-out warp seeds, Sutura fits torn-tissue correspondence to a median 99 → 106 px (5-seed). Three caveats: (i) supervised-on-target versus cold unsupervised comparison; (ii) soft regression that is ∼138 px under hard assignment; (iii) flat severity curve partly structural. The model is a proof-of-concept showing that a graph cross-attention architecture can fit torn-tissue correspondence at near sub-pitch resolution, not a deployable tool.

### 5.3 The diagnosed generalisation failure

Under LODO, Sutura is ∼1.8–3.6× worse than PASTE2 on unseen tissue, flat across severity. The mechanism is diagnosed: cross-donor batch shift pushes held-out expression off-distribution; the encoder produces non-discriminative embeddings; cross-attention collapses toward uniform; predictions regress to the tissue centroid. This is an instance of a recognised hard failure mode of deep ST models, quantified here with five-seed CIs and traced end-to-end to its cause.

### 5.4 Contrastive correspondence loss

A targeted InfoNCE-style contrastive loss (single-seed, all three LODO folds) closes the gap substantially on two donors: best held-out median falls to 816 → 949 px (S1) and 749 → 826 px (S2), reaching ∼1.1 – 1.2× of PASTE2 at worst-case tear. The fix helps only modestly on S3 ( ∼13%, still ∼2.4 × PASTE2) and does not surpass PASTE2 on any donor. Donor-invariance is improved, not solved.

### 5.5 Limitations

(1) Three donors: *n* = 3 donors is the true replication unit for cross-donor claims. (2) Single dataset and platform: all results on spatial-LIBD DLPFC 10x Visium. (3) Synthetic, rigid tears: real tears can fragment tissue and alter local expression. (4) Approximate ground truth: ∼8 px residual floor, one-pitch hard-assignment floor. (5) No independent learned baseline with code available: STaCker and INST-Align could not be executed. (6) Model-fixed CIs: warp-realisation variance only. (7) Contrastive runs are single-seed.

### 5.6 Future directions

(1) Batch integration before SVD (Harmony, Scanorama) so heldout donors project in-distribution. (2) Multi-donor encoders trained to learn donor-invariance directly. (3) Benchmarking one learned-deformation method with released code on this exact tear benchmark, to establish whether non-generalisation is Sutura-specific or field-wide. (4) A second tissue and platform. (5) Evaluation on a real torn section.

## Data and Code Availability

All code is available at https://github.com/Sutura-Genomics/sutura-paper, including model implementation, training and evaluation scripts, baseline wrappers (PASTE2, STalign, GPSA), pretrained checkpoints, all result CSVs, figure generation scripts, and a quick-start inference notebook. The spatialLIBD DLPFC Visium dataset is publicly available from the CellCharter benchmark distribution (Figshare article 22004273, CC BY 4.0).

## Author Contributions

R.M. designed the model architecture, ran the multi-seed PASTE2/STalign/GPSA benchmarks, the in-sample Sutura experiments, the leave-one-donor-out experiments, and the contrastive correspondence loss matrix, and generated all figures. S.L. wrote the Methods, Related Work, and Discussion sections, contributed to experimental design, and curated the cross-donor evaluation pipeline. S.S.L. supervised the project, contributed to experimental design, provided clinical and computational guidance on tumour tissue analysis, and reviewed the manuscript. All authors approved the final version.

## Acknowledgments

We thank the spatialLIBD team at the Lieber Institute for Brain Development for making the DLPFC Visium dataset publicly available, and the developers of PASTE2 (Raphael Lab), STalign (Fan Lab), and GPSA (Engelhardt Lab) for open-source code that enabled this benchmark.

